# Indolent primary cutaneous B-cell lymphomas resemble persistent antigen reactions without signs of dedifferentiation

**DOI:** 10.1101/2022.12.16.520801

**Authors:** Johannes Griss, Sabina Gansberger, Inigo Oyarzun, Mathias C. Drach, Vy Nguyen, Lisa E. Shaw, Ulrike Mann, Stefanie Porkert, Matthias Farlik, Wolfgang Weninger, Werner Dolak, Bertram Aschenbrenner, Beate M. Lichtenberger, Christine Wagner, Ingrid Simonitsch-Klupp, Stephan N. Wagner, Constanze Jonak, Patrick M. Brunner

## Abstract

Primary cutaneous B-cell lymphomas comprise a heterogeneous group of extranodal non-Hodgkin lymphomas. While primary cutaneous diffuse large B-cell lymphoma leg type (pcDLBCL-LT) is highly aggressive, the two other subtypes, primary cutaneous follicle centre lymphoma (pcFCL) and primary cutaneous marginal zone lymphoma (pcMZL), usually follow an indolent course.

To better understand the molecular landscape of these entities, we performed single-cell RNA sequencing of pcFCL, pcMZL, and pcDLBCL-LT skin lesions and compared them to B-cell rich lymphoid proliferation (rB-LP) lesions, gastric MALT lymphoma, nodal FCL, and nodal DLBCL.

Our data show that these lymphomas can be clearly distinguished from each other on a transcriptomic level based on scRNA-seq. pcMZL, pcFCL, and rB-LP all exhibited a persistent germinal centre reaction as evidenced by the presence of required support cells and continuous somatic hypermutation within the expanded clone. By contrast, malignant clones of pcDLBCL-LT and gastric MALT lymphoma lesions lacked these features. Further, pcMZL top expanded clones were developing within lesions from naïve and not post-germinal centre B cells as currently presumed. Therefore, pcMZL may represent a non-malignant reaction against a yet to be determined antigen. Conversely, in pcFCL, B cells showed a larger amount of clonal expansion. The lack of further differentiation of these B cells may explain its indolent clinical course. In contrast to pcDLCBL-LT, our data thus indicate that pcMZL and pcFCL, similar to rB-LP are characterised by a functional germinal centre reaction likely driven by (a yet unknown) antigen recognition, which supports the classification of pcMZL as a lymphoproliferative disease.

**Key points:**

- Indolent B cell neoplasms are uniquely characterised by ongoing, antigen-driven germinal centre reactions.
- Primary cutaneous B cell lymphomas are distinct entities compared to their systemic counterparts.

## Introduction

Primary cutaneous B-cell lymphomas comprise a heterogeneous group of extranodal non-Hodgkin lymphomas^1^. The most common subtypes include primary cutaneous marginal zone lymphoma (pcMZL), primary cutaneous follicle centre lymphoma (pcFCL), and primary cutaneous diffuse large B-cell lymphoma, leg type (pcDLBCL-LT)^1^. PcMZL and pcFCL are generally indolent conditions with a 5-year disease-specific survival of >95%^2^. In contrast, pcDLBCL-LT is an aggressive disease with 5-year survival rates between 20 - 60%.^3^

For pcMZL, this clinical observation led to a reclassification to a lymphoproliferative disorder in the International Consensus Classification (ICC) joint classification referring to their class-switched cases only^2^. These comprise IgG or IgA positive B-cell proliferations, which follow an indolent clinical course. PcMZL may be preceded by reactive B-cell rich lymphoid proliferations (rB-LP, formerly called “pseudolymphoma”), matching that pcMZL are not veritable lymphomas^4^. In line, several studies showed that only around 5% of B cells within pcMZL lesions correspond to a top expanded clone ^5,6^. Nevertheless, the WHO classification currently maintains the term lymphoma for pcMZL leading to a conflict between these two classifications^7^.

Previous gene expression analyses identified characteristic B-cell phenotypes for each primary cutaneous B-cell lymphoma subtype. pcMZL B cells were found to be most similar to plasma cells and pcFCL showed a more germinal centre (GC)-like phenotype, whereas pcDLBCL-LT lesions were most consistent with activated B cells.^8–10^ In cutaneous B-cell lymphomas, clonality was not always detectable using routine molecular methods, or when a surrogate such as light-chain restriction was used.^11^ In those cases, the differentiation from rB-LP can be difficult.^12^ To distinguish pcMZL from rB-LP, skin flow cytometry was found to have superior sensitivity than immunohistochemistry, in situ hybridization or immunoglobulin heavy chain gene rearrangement examinations.^12^ However, its routine use is difficult as it requires fresh tissue, is prone to technical challenges during processing, and is not widely available.^12^ Apart from a more precise molecular characterisation of the disease, the terminology of lymphoma versus lymphoproliferative disorder or cutaneous lymphoid hyperplasia impacts patient education and especially the patients’ perception of their disease^13,14^.

Here, we present a single-cell RNA sequencing characterisation of primary cutaneous B-cell lymphomas and rB-LP and compare these findings to data from their non-cutaneous B-cell counterparts.

## Methods

### Patient recruitment and sample processing

Patients were recruited at the Medical University of Vienna, Austria. Skin punch biopsies or biopsies during gastroscopy were taken after obtaining written informed consent, under a protocol approved by the Ethics Committee of the Medical University of Vienna (EK 1360/2018). For all patients with rB-LP, borrelia infection was ruled out. After processing using the Skin Dissociation Kit by Miltenyi Biotech (Bergisch Gladbach, Germany), ^15–17^ single-cell suspensions were subjected to scRNA-seq using the Chromium Single Cell Controller and Single Cell 5’ Library & Gel Bead Kit v2 (10X Genomics, Pleasanton, CA) according to the manufacturer’s protocol. B cell receptor (BCR) sequences were enriched from cDNA using the VDJ Kit workflow by 10X Genomics. Sequencing was performed using the Illumina NovaSeq platform and the 150bp paired-end configuration.

### Data analysis

Raw data from scRNA-seq was preprocessed using Cell Ranger version 7.0.1 invoking the command “multi” and aligned to the human reference genome assembly “refdata-gex-GRCh38-2020-A” and Cell Ranger V(D)J segment reference “refdata-cellranger-vdj-GRCh38-alts-ensembl-7.0.0”.

IgBLAST^18^ v1.20 was used on the BCR filtered contigs to quantify the somatic hypermutation (SHM), presented as *”1 - identity in variable region”*.

B cell clones were identified from the BCR data through the following steps. Cells featuring more than two contigs or multiple contigs for the same BCR chain (heavy or light chain), and clones with more than two CDR3 chains, were removed. Subsequently, if one of the two most frequent clones only contained one heavy or one light chain were merged with clones were both chains were identified if all cells shared the same V, C, and J genes as the evaluated clonotype, but only if all cells displayed at least 85% sequence identity in both light and heavy CDR3 sequences compared to the reference sequence of the evaluated clonotype, using global pairwise alignment from Biostrings^19^ R package v2.68 (BLOSUM100 matrix with a gap opening and gap extension penalty values of 10 and 4 respectively).

Every sample was processed using R (version 4.1.0) and Seurat (version 5.0.1)^20^. Ambient RNA was removed using DecontX^21^ and doublets removed using scDblFinder^22^. During the quality assessment, only cells with at least 500 genes and a maximum of 10%, 5% and 1% of mitochondrial, haemoglobin and platelet genes respectively were kept for downstream analysis.

Afterwards, samples were normalised and scaled regressing for percentage of mitochondrial genes and sample origin. 30 dimensions were used when invoking the Seurat functions RunPCA, FindNeighbors and RunUMAP. Subsequently, samples were integrated using the FastMNN method^23^. Leiden algorithm was applied on the shared nearest neighbour graph clustering prior to cell annotation as suggested by Heumos, L. *et al.*^24^. In order to achieve a deeper annotation, B and T cells were subclustered, variance was stabilised using SCTransform v2^25^ and ScaleData respectively with 15 and 20 dimensions respectively for subclusterings.

### Immunohistochemistry

Multiplex immunostainings were conducted as previously described^26^. Briefly, 4μm sections were deparaffinized and antigen retrieval was performed in heated citrate buffer (pH 6.0) and/or Tris-EDTA buffer (pH 9) for 30 min. Thereafter, sections were fixed with 7.5% neutralised formaldehyde (SAV Liquid Production GmbH, Flintsbach am Inn, Germany). Each section was subjected to 6 successive rounds of antibody staining, consisting of protein blocking with 20% normal goat serum (Dako, Glostrup, Denmark) in PBS, incubation with primary antibodies, biotinylated anti-mouse/rabbit secondary antibodies and Streptavidin-HRP (Dako), followed by TSA visualisation with fluorophores Opal 520, Opal 540, Opal 570, Opal 620, Opal 650, and Opal 690 (PerkinElmer, Waltham, MA, USA) diluted in 1X Plus Amplification Diluent (PerkinElmer), Ab-TSA complex-stripping in heated citrate buffer (pH 6.0) and/or Tris-EDTA buffer (pH 9) for 30 minutes, and fixation with 7.5% neutralised formaldehyde. Thereafter, nuclei were counterstained with DAPI (PerkinElmer), and sections were mounted with PermaFluor fluorescence mounting medium (Thermo Fisher Scientific, Waltham, MA, USA). Multiplexed slides were scanned on a Vectra Multispectral Imaging System version 2 following the manufacturer’s protocol (InForm 2.4, Perkin Elmer). All phenotyping and subsequent quantifications were performed blinded to the sample identity.

### Data availability

Gene level expression and VDJ sequencing data for new samples are available on GEO (GSE218861). Data from healthy control samples are available under GSE173205^27^.

## Results

### scRNA-seq based characterisation matches the current histopathological understanding of primary cutaneous B-cell lymphomas

We obtained fresh biopsies from patients diagnosed with pcMZL (n = 9), pcFCL (n = 5), pcDLBCL-LT (n = 4), or cutaneous rB-LP (n = 5), and normal skin from healthy volunteers (NHS, n = 4) (Supplementary Table 1). Samples were analysed using single-cell RNA-sequencing (scRNA-seq) coupled with B-cell receptor (BCR) sequencing. After quality control, we obtained data for 268.224 cells (pcDLBCL-LT: 16.908, pcFCL: 71.720, pcMZL: 109.028, rB-LP: 51.336, NHS: 19.232).

Using canonical markers (Supplementary Figure 1), we were able to identify T cells and NK cells, B cells, plasma cells, blood vessel and lymphatic endothelial cells, dendritic cells, plasmacytoid dendritic cells, macrophages, melanocytes, keratinocytes, smooth muscle cells, and fibroblasts (Figure 1A, Supplementary Figure 2).

**Figure 1.**
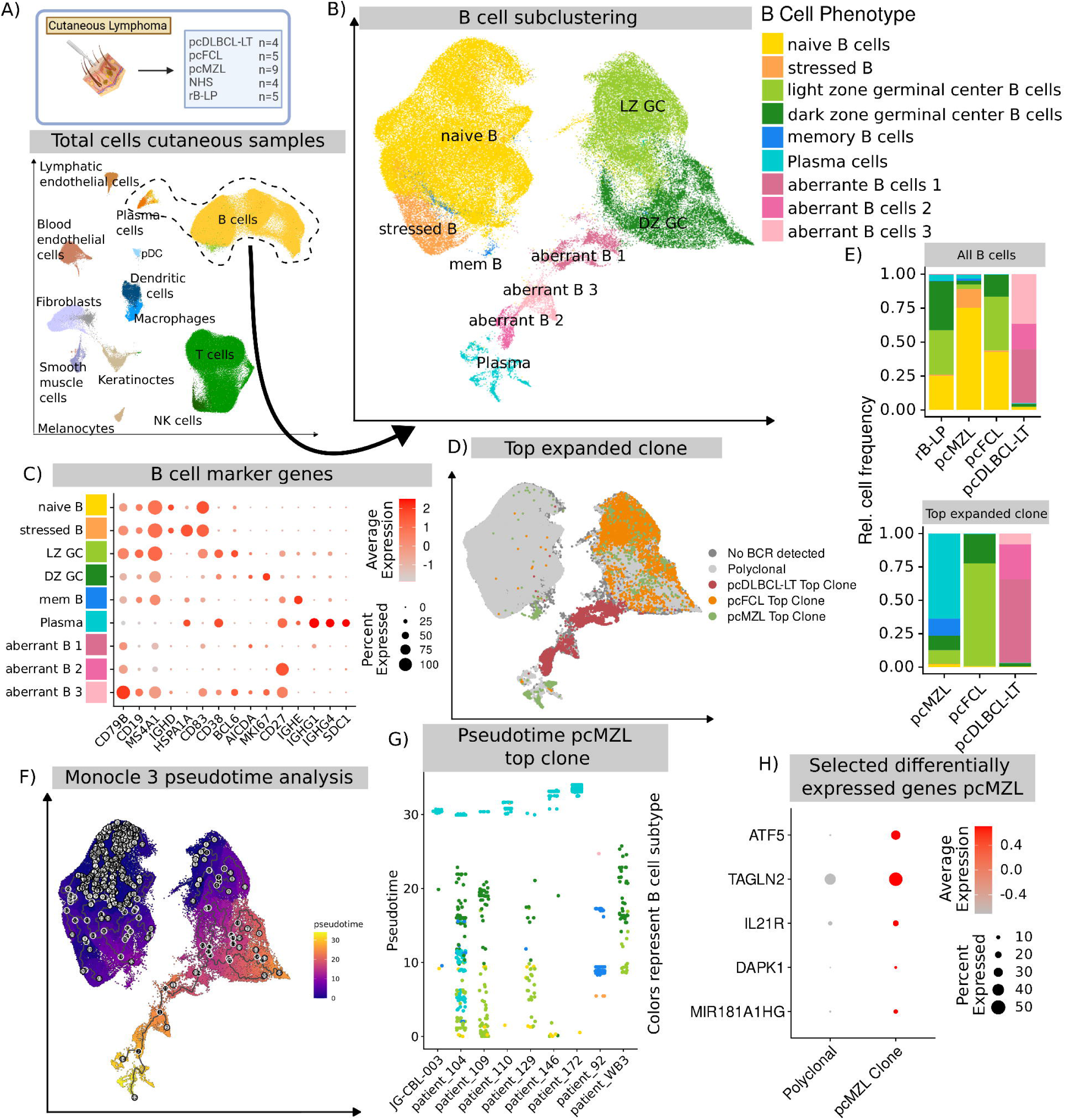
**A)** Schematic overview of all cutaneous samples and UMAP embedding of the respective scRNA-seq data. **B)** UMAP embedding of the subclustered B cells (including B, and Plasma cells) of all cutaneous samples. **C)** Dot plot of key B cell markers used to identify respective B-cell subtypes. Colour intensity represents average expression and point size the fraction of expressing cells. **D)** UMAP embedding of the subclustered B cells highlighting the results of the BCR receptor sequencing. Colours represent the top-expanded clone of all samples per disease. **E)** Frequency of B-cell subtypes per disease shown for all B cells (top) and only B cells part of the top-expanded clone. **F)** UMAP embedding of the B cell clubclustering highlighting the results of the monocle3-based pseudotime analysis. Black lines represent the identified trajectories. **G)** Pseudotime of each individual cell from the top-expanded clone in pcMZL per sample. Colours represent the identified B-cell subtypes. **H)** Dot plot of differentially expressed genes between clonally expanded and polyclonal B cells in pcMZL. Dot size represents the number of cells expressing the respective genes, colour intensity represents the abundance value.

We further subclustered all B cells in order to arrive at a detailed phenotypic characterisation (Figure 1B). This revealed naïve B cells, as identified by CD19, MS4A1 (CD20), and IGHD, germinal centre (GC) B cells, as identified by CD38, AICDA, and BCL6 that were further subdivided into a light zone (LZ) and dark zone (DZ) GC B-cell population, based on the presence or absence of MKI67 (Figure 1C). Memory B cells were characterised by their expression of CD19, MS4A1 (CD20), and CD27 with the absence of CD38 and BCL6 (Figure 1C). Plasma cells were identified through the expression of CD27, CD38, SDC1 (CD138), and the lack of MS4A1 (CD20) (Figure 1C). This overall composition could be confirmed using a 6-plex based immunohistochemistry analysis of archival tissue samples from patients diagnosed with rB-LP, pcMZL, and pcDLBCL-LT (n = 19, Supplementary Figure 3). Thus, we were able to identify the full spectrum of canonical B-cell phenotypes ranging from naïve B cells to plasma cells.

In addition, we observed aberrant B-cell populations, all originating from pcDLBCL-LT, that formed distinct clusters and could not be classified using canonical B-cell markers (Figure 1B-E). These were in part MS4A1 (CD20)+, but also expressed MKI67 without the typical expression of CD38 and CD27 for canonical germinal centre B cells (“aberrant B1”, Figure 1C). Additionally, we observed MS4A1 (CD20) negative B cells that lacked the expression of SDC1 (CD138) and high levels of immunoglobulin associated genes that would be expected for plasma cells (“aberrant B2”, Figure 1C). This indicates the pcDLBCL-LT contains B cells that no longer match the physiological B-cell phenotypes.

### pcMZL originates from pre-germinal centre naïve B cells and follows the canonical B-cell differentiation trajectory

Through our BCR sequencing data, we were able to clearly attribute B cells to the top expanded clone in each disease (Figure 1D). In each sample, we were thus able to unambiguously distinguish the clonally expanded (presumed malignant) B cells from the polyclonal bystander infiltrate (Supplementary Material 1).

Matching current classifications^28^, pcMZL and rB-LP uniquely contained relevant numbers of plasma cells (Figure 1E). Conversely, pcFCL samples showed a dominance of LZ GC B cells (Figure 1E). pcDLBCL-LT uniquely contained aberrant B-cell phenotypes and lacked relevant numbers of naïve B and canonical plasma cells (Figure 1E). Our data is therefore in-line with the current histopathological understanding with distinct phenotypic compositions of the B-cell infiltrate for each entity.

pcMZL top clones are considered to derive from post GC B cells ^28,29^. However, we found that clonally expanded B cells from our pcMZL samples spanned all canonical peripheral B-cell phenotypes, from naïve B cells up to terminally differentiated plasma cells (Figure 1E). We used monocle3 to perform a pseudotime analysis of this data, which matched the canonical B cell differentiation (Figure 1F, Supplementary Figure 4). An analysis of the clonally expanded B cells of each pcMZL sample highlighted that the clone seemed to differentiate in each individual sample (Figure 1G). This was consistent regardless of whether the sample exhibited a predominance of plasma cells or germinal centre B cells and was not limited to specific subtypes of pcMZL as previously described^29^. This suggests that pcMZL clones develop from naïve, pre-GC B cells irrespective of the phenotypic composition.

To assess potential functional differences, we performed a differential expression analysis between clonally expanded and polyclonal B cells in pcMZL (Figure 1H, Supplementary Figure 5, Supplementary Material 2). We observed an up-regulation of genes both inducing and inhibiting proliferation in clonally expanded plasma cells. ATF5, for example, is a known pro-survival transcription factor^30^ and IL21R is directly responsible for the IL21 driven expansion of plasma cells^31^. In contrast, MIR181A1HG inhibits NFkB signalling and thereby B-cell proliferation^32^ and DAPK1 induces apoptosis and is downregulated in chronic lymphocytic leukemia^33^. Finally, the activation marker TAGLN2^34^ was up-regulated in clonally expanded plasma cells. In clonally expanded germinal centre B cells, we primarily observed a down-regulation of MHC-II associated genes. Other differences were mainly driven by single samples. Overall, we observed differences that are compatible with activated B cells, where the respective anti-proliferative regulatory mechanisms seem to be still intact.

### Non-class switched pcMZL samples show aberrant phenotypes

Several authors currently differentiate between a class switched and non-class switched variant of pcMZL, where only the non-class switched variant is considered a true lymphoma^2,28^. In our series, one sample (patient 92) had a clear IgM+ phenotype among the clonally expanded B cells (Figure 2A). When processing these B cells individually, we observed that the clonally expanded B cells formed a distinct cluster from the polyclonal ones (Figure 2B). This cluster was MS4A1 (CD20)-, CD27+, IGHM+, with a MKI67+ subpart (Figure 2B) that showed low expression of CD38 and AICDA but no relevant expression of IGHG1-4- (Supplementary Figure 6). There were no B cells in this sample that expressed CD138 (SDC1), which would be expected for CD20- B cells (Supplementary Figure 6). While many aspects of these cells, such as proliferation and potential somatic hypermutation (expression of AICDA) is reminiscent of germinal centre B cells, the lack of CD20 is incompatible with this phenotype, matching reports of CD20 loss in systemic B cell lymphoma^35^. Clinically, patient 92 has shown an indolent course with two cutaneous recurrences and no sign of systemic disease within a total follow up of 12 years. Therefore, our data suggests that this sample’s clone does consist of aberrant, non-canonical B cells yet with no apparent influence on this patient’s clinical course.

**Figure 2.**
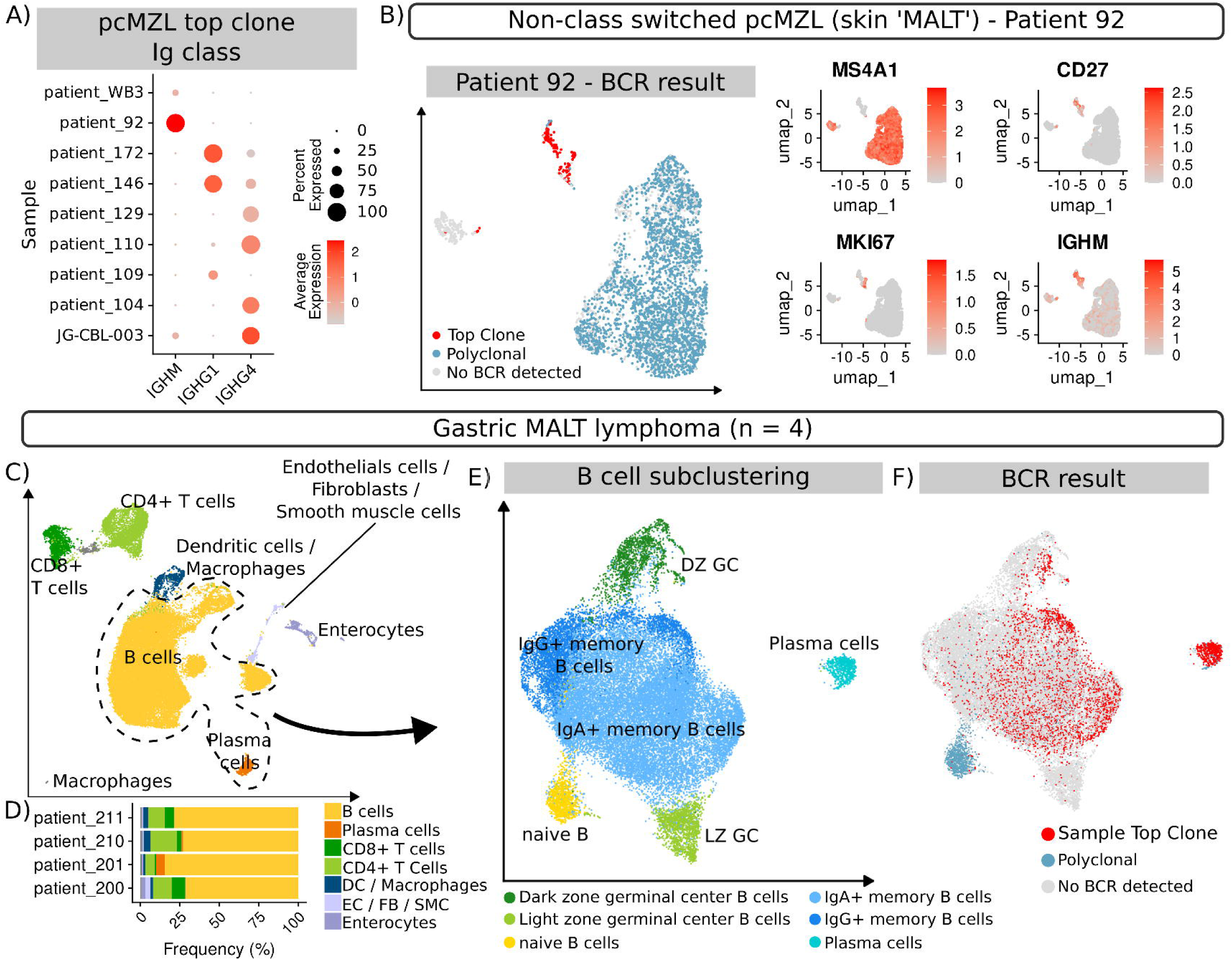
**A)** Dot plot showing the abundance of immunoglobulin isotype genes in the top expanded clone per pcMZL sample. **B)** UMAP embedding of a subclustering of the B cells of sample 92. Colours in the left panel represent the result of the BCR sequencing and in the right panels the abundance of selected marker genes. **C)** UMAP embedding of all cells characterised from the four MALT samples. The dotted line highlights the B cells used for the subclustering. **D)** Relative abundance per cell type and sample of the MALT samples. **E, F)** UMAP embedding of the subclustering of all B and plasma cells from the MALT samples highlighting the **E)** identified B-cell subtypes and **F)** the results of the BCR sequencing.

A second sample, WB3, also showed a non-class switched clone (Figure 2A). Nevertheless, these clonally expanded cells matched canonical germinal centre B cells (Supplementary Figure 7). In this sample, plasma cells were all polyclonal and IgG4 positive (Supplementary Figure 7). In a routine histopathological assessment, sample WB3 would therefore be classified as a class-switched pcMZL. This observation highlights that in this specific sample, the histopathological differentiation of class switched *vs.* non-class switched pcMZL is likely not yet precise enough to distinguish clinically relevant subtypes.

### pcMZL is distinct from gastric MALT lymphoma

Gastric mucosa associated lymphoid tissue (MALT) lymphoma is one of the most frequent marginal zone lymphomas^36^, but whether it is similar to pcMZL is unclear. We therefore acquired biopsies from 4 patients diagnosed with gastric MALT lymphomas refractory to *H. pylori* eradication (Supplementary Table 1). Based on canonical markers, we were able to identify B and plasma cells, both CD4+ and CD8+ T cells, a cluster of dendritic cells and macrophages, enterocytes, and a cluster of epithelial cells, smooth muscle cells, and fibroblasts (Figure 2C). In contrast to pcMZL samples (Supplementary Figure 2), all MALT lymphoma were dominated by B cells, which represented around 75% of all cells (Figure 2D). These B cells primarily consisted of CD27+ memory B cells which all showed a class-switch to either IgG or IgA (Figure 2E, Supplementary Figure 8). The top expanded clone was dominated by memory B and plasma cells, with lower numbers of germinal centre B cells. Naïve B cells were polyclonal (Figure 2F). This indicates that gastric MALT lymphoma develops within a germinal centre reaction. Overall, these data show that pcMZL is a distinct entity from gastric MALT lymphoma, both with regard to the clonal expansion, as well as the composition of the B-cell infiltrate and the presumed cell of origin.

### Clonally expanded B cells in pcFCL are uniquely confined to germinal centre B cells

pcFCL is histopathologically differentiated from pcMZL primarily through the expression of BCL6^28^, which can be challenging for cases of pcFCL with plasmacytic differentiation^37,38^. In our pcFCL samples, the top expanded clone was strictly limited to GC B cells, while B-cell phenotypes other than GC cells only contained polyclonal B cells (Figure 1E). This was consistent with a published pcFCL dataset that we re-analsed, where clonally expanded B cells were limited to GC B-cell clusters^39^ (Figure 3 A, B). To assess whether this is true for all follicle centre lymphomas, we re-processed public data from a study characterising 20 samples from patients diagnosed with sFCL^40^ (Figure 3C). Overall, we did not observe any aberrant B-cell phenotypes in these samples (Supplementary Figure 9). While the samples and the top expanded clone were dominated by germinal centre B cells, the clone also contained memory B and plasma cells (Figure 3D). Only naïve B cells consisted solely of polyclonal B cells. This highlights that the confinement of the clone to one B-cell phenotype is a unique feature of pcFCL. This enables an unambiguous classification based on scRNA-seq data.

**Figure 3.**
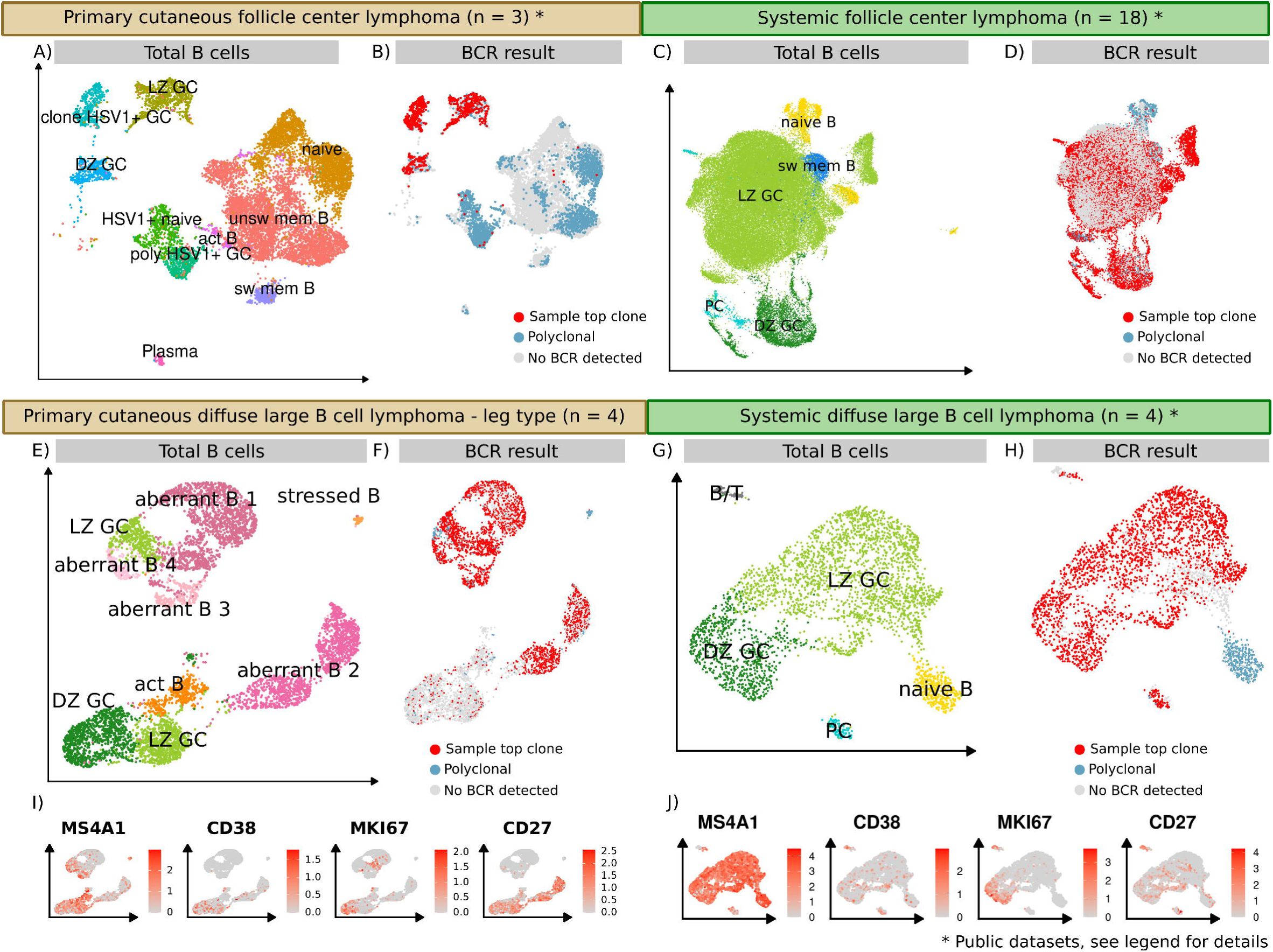
**A**, **B)** UMAP embedding of the subclustering of the B cells from a public pcFCL sample from Ramelyte *et al.*^39^. Colours indicate the identified B-cell phenotype with HSV+ cells infected with the investigated oncolytic virus (**A**). **C, D)** UMAP embedding of the subclustering of the B cells from public sFCL samples from Han *et al.*^40^. Colours indicate identified B-cell subtypes (**C**). **E, F)** UMAP embedding of the subclustering of the B cells from the pcDLBCL-LT samples. **G, C)** UMAP embedding of the subclustering of B cells from sDLBCL samples from Steen *et al.*^41^. **B, D, F, H** represent the respective results of the BCR sequencing. **I, J)** Expression of key B-cell markers in the (**I**) pcDLCBL-LT and (**J**) sDLBCL B cells.

### pcDLBCL-LT is characterised by aberrant B-cell phenotypes

Clonally expanded B cells in pcDLBCL-LT could not be annotated following the canonical B-cell development in our samples (Figure 1C). A detailed analysis of only pcDLBCL-LT B cells confirmed these initially observed aberrant phenotypes (Figure 3E, F). We observed a loss of MS4A1 (CD20) without the expression of SDC1 (CD138) or CD38 as expected of CD20- plasma cells (Figure 3I, aberrant B1). There were two clusters of proliferating (MKI67+) B cells. While one matched DZ GC B cells, the other was MS4A1 (CD20)-, with no expression of CD38 or CD27 required for germinal centre B cells (Figure 3I, aberrant B2). These findings were consistent with publicly available scRNA-seq data^39^ (re-analyzed in Supplementary Figure 10). To test whether this is typical of all DLBCL lymphomas, we reprocessed a public dataset with 4 samples of sDLBCL^41^ (Figure 3 G, H). In contrast, clonally expanded B cells from sDLBCL samples showed canonical marker profiles with no loss of MS4A1 (CD20) (Figure 3J). Additionally, the clonally expanded B cells also contained canonical plasma cells (Figure 3H). Therefore, pcDLBCL-LT was the only lymphoma in our analysis where all samples consisted of non-canonical B cell phenotypes.

### pcMZL shows significantly lower clonal expansion than other cutaneous and systemic lymphomas

Clonal expansion is considered a hallmark of lymphomas and can be tracked in B cells through BCR sequencing^42^. Using our BCR data, we were able to accurately quantify the level of clonal expansion in each sample (Figure 4A). In pcMZL and rB-LP, clonally expanded B cells represented a median of 4% and 2% respectively of all B cells, matching previous reports^5^. Outliers observed were likely due to sequencing artefacts (Supplementary Figure 11, Supplementary Figure 12). This was in contrast to pcFCL and pcDLCBL-LT which showed median clonal expansions of 52% and 97% respectively. Similarly, in gastric MALT lymphoma the expanded clone accounted for a median of 77% of the total B-cell infiltrate. Moreover, sFCL and sDLBCL were similarly dominated by a single expanded clone, with a median expansion of 96% and 85% respectively. Overall, the rate of clonal expansion substantiates the theory that pcMZL is more similar to a lymphoproliferative disorder than a true lymphoma.

**Figure 4.**
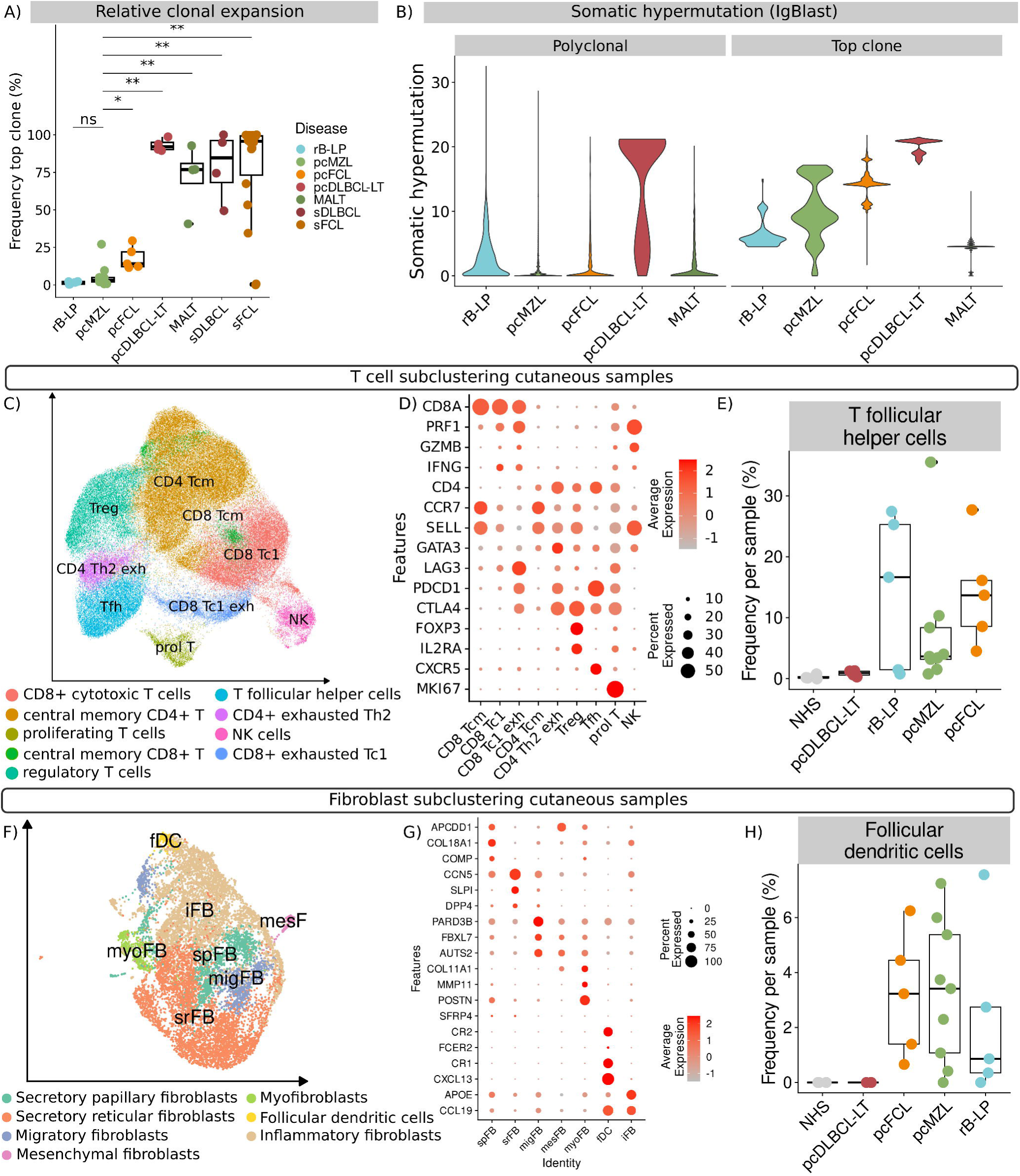
**A)** Relative proportion of the top expanded clone per sample and disease. Values are normalised based on the total number of B cells. Stars represent significant levels based on a Wilcoxon rank sum test with Bonferroni correction (* < 0.01, ** < 0.05, n.s. > 0.05). **B)** Somatic hypermutation in the top expanded B-cell clone and polyclonal B cells per disease as derived through the identity mapping of IgBLAST. Somatic hypermutation is quantified through the difference to the canonical sequence. Data is shown separately for the top expanded clone and the polyclonal infiltrate. **C)** UMAP embedding of a subclustering of the T cells from all cutaneous samples. **D)** Canonical markers used to identify key T cell subtypes. Size of the dots represent the percentage of cells expressing the marker, colour intensity represents the average expression. **E)** Relative proportion of T follicular helper cells among all identified T cells per sample (coloured points). Lower and upper hinges of the boxplots correspond to the first and third quartiles, whiskers extend to a maximum of 1.5 of the interquartile range and represent the minimum and maximum value within this range. **F)** UMAP embedding of a subclustering of all fibroblasts from the cutaneous samples. **G)** Dot plot showing canonical markers used to identify key fibroblast subsets. **H)** Relative proportion of follicular dendritic cells (fDC) normalised based on all identified fibroblasts per sample (coloured points). Lower and upper hinges of the boxplots correspond to the first and third quartiles, whiskers extend to a maximum of 1.5 of the interquartile range and represent the minimum and maximum value within this range.

### Indolent B-cell lymphomas and rB-LP are characterised by ongoing germinal centre reactions

The germinal centre reaction with its ongoing somatic hypermutation is a key part of B-cell function. We used IgBLAST^18^ to quantify the rate of somatic hypermutation in all cutaneous samples (Figure 4B). In all diseases, the top expanded clone showed a higher rate of somatic hypermutation compared to the polyclonal B cells. In pcMZL and rB-LP, the most expanded clone had the widest range of mutations. This matches the assumption that these B cells entered the germinal centre reaction and subsequently acquired mutations to improve antigen specificity, which is in line with the results of our pseudotime analysis of pcMZL samples (Figure 1G). pcFCL similarly showed increasing rates of somatic hypermutation, yet starting at a considerably higher level (Figure 4B). This matches our observation that the clone in pcFCL did not contain any naïve B cells (Figure 1E). Yet, there is evidence of an ongoing acquisition of mutations. Clonally expanded B cells in pcDLCBL-LT showed the highest rates of somatic hypermutation but spanning a much smaller range (Figure 4B). Despite our characterization of germinal centre B cells in pcDLBCL-LT samples, these data suggest that these cells are largely no longer acquiring new mutations typically associated with the physiological germinal centre reaction. This finding further supports the aberrant phenotypes observed in these samples. In MALT lymphoma samples, polyclonal B cells showed a similar rate of somatic hypermutation as the polyclonal B cells in pcMZL and pcFCL. Yet, the top expanded clone centred around a single value of somatic hypermutations. This matches our phenotypic data where clonally expanded B cells in MALT samples primarily consisted of post-germinal memory B and plasma cells that no longer undergo the germinal centre reaction (Figure 2E, F). This data suggests that the three indolent cutaneous B-cell reactions, rB-LP, pcMZL, and pcFCL uniquely retain an intact germinal centre reaction.

The germinal centre reaction depends on several other cell types, most importantly CXCR5+ T follicular helper cells (Tfh) and follicular dendritic cells (fDC)^43^. We therefore subclustered the T cells from all cutaneous samples (Figure 4C). We were able to identify CD8+, PRF1+, IFNG+ Tc1 polarised cells (CD8 Tc1), some of which were LAG3+ as a sign of exhaustion (CD8 Tc1 exh, Figure 4D). Further, we observed CD4+, CCR7+, SELL+ CD4 central memory T cells (Tcm), FOXP3+, IL2RA+ regulatory T cells (Treg), MKI67+ proliferating T cells (prol T), PDCD1+, CTLA4+ exhausted CD4+ T cells (CD4 Th2 exh), as well as CXCR5+ T follicular helper cells (Tfh). Tfh were only present in rB-LP, pcMZL, and pcFCL samples (Figure 4E), matching previous reports where Tfh were histologically observed in all cases of pcMZL^44^.

Matching their mesenchymal origin with an assumed fibroblast-like progenitor^45^, fDC are part of fibroblast clusters in scRNA-seq data. We therefore further subclustered all fibroblasts from the cutaneous samples and identified APCDD1+, COL18A1+ secretory papillary (spFB), CCN5+, SLPI+, DPP4+ secretory reticular (srFB), PARD3B+, FBXL7+, AUTS2+ presumably migratory (migFB), COL11A1+, MMP11+, POSTN+ mesenchymal (mesFB), SFRP4+ myo (myoFB), APOE+CCL19+ inflammatory (iFB) fibroblasts, CR2+, FCER2+, CR1+ CXCL13+ follicular dendritic cells (fDC) and S100B+SOX10+ Schwann cells (SC) (Figure 4F, G). Similarly to Tfh, fDC were only observed in rB-LP, pcMZL, and pcFCL samples (Figure 4H). Therefore, with the detection of an ongoing somatic hypermutation and the presence of Tfh and fDC, samples of rB-LP, pcMZL, and pcFCL uniquely contained all required aspects of a functional germinal centre reaction.

## Discussion

The indolent behaviour of pcMZL, pcFCL, and rB-LP has sparked discussions on their actual nature, and led to conflicting classifications with respect to pcMZL^7^. Previous studies showed that patients’ quality of life is directly influenced by whether a disease is classified as a lymphoma or not^14^. This highlights that we need to arrive at a clear statement concerning the nature of these entities for patients, as well as for clinical practice.

Our data presents consistent evidence that the investigated three indolent B-cell diseases, rB-LP, pcMZL, and pcFCL are uniquely characterised by ongoing germinal centre reactions. We were able to detect both continuous somatic hypermutation, as well as the required support cells in these samples. Germinal centre reactions are generally considered a sign of the formation of tertiary lymphoid structures^46^, are primarily observed in cancer^47^ and autoimmune diseases^48^ and are a sign of chronic, antigen-specific immune responses^49^. This matches multiple reports where both pcMZL and pcFCL were linked to infections^50–52^ and responded to antibiotic therapies or vaccinations^53–55^. Next to the ongoing germinal centre reaction, the top expanded clones in pcMZL represented only a low fraction of the overall B cells, matching previous reports^5^. This is in contrast to all other lymphomas, including MALT and matches reports that (systemic) DLBCL is driven by antigen-independent B-cell receptor activation^56^. Additionally, these B cells followed physiologic B-cell trajectories, originating from lesional naïve B cells, counter to current hypotheses and in contrast to MALT and pcDLBCL-LT^28^. This further matches previous studies that were unable to find consistent driver mutations in pcMZL^28^. Overall, pcMZL therefore does not show any of the characteristics found in the other investigated B-cell lymphomas but matches rB-LP with the sole difference that pcMZL developed a presumably antigen-directed, expanded clone. We therefore, for the first time, present molecular evidence that aligns with the concept that pcMZL is a lymphoproliferative disorder and not a true lymphoma.

Our data further questions the proposed class-switch based classification of pcMZL into a lymphoproliferative disorder or a lymphoma^4^. In our data, one non-class switched sample may contain aberrant B cells, yet this patient showed an indolent course within a follow up of 12 years. In the second case, the top expanded clone consisted mainly of non-class switched, canonical germinal centre B cells. Yet, the more abundant polyclonal plasma cells were all class switched. In routine histopathological assessments, this sample would therefore have been classified as class switched. This highlights that the proposed practice of defining class switched and non-class switched pcMZL is not necessarily linked with a high risk of systemic disease.

In contrast to pcMZL, pcFCL showed a comparably high level of clonal expansion. Yet, these clones were still undergoing somatic hypermutation as part of the germinal centre reaction. This is in contrast to a series of pcFCL cases where the authors could not detect signs of somatic hypermutation^57^. It is unclear whether this is a result of the higher sensitivity of our scRNA-seq data or the presence of different subsets of pcFCL. In contrast to sFCL, our samples did not show any sign of further differentiation but appeared to remain contained within a single phenotype. Previous studies did identify putative driver mutations in pcFCL^58^. We believe that these mutations were acquired during the ongoing germinal centre reaction, which is believed to increase the risk for such mutations^59^. Thereby, the clone may be able to expand, yet still requires the support from the germinal centre environment. This locks it in its single phenotype and may explain pcFCL’s indolent behaviour.

A question we currently cannot address is whether FCL primarily occurring in the skin with bone marrow involvement is different to pcFCL. Senff *et al.* showed that 4.5% of patients with FCL primarily occurring in the skin had bone marrow involvement without other signs of systemic disease^60^. In their study, these nine patients showed significantly shorter survival times compared to patients with pcFCL. While in our patients CT studies showed no sign of systemic disease, bone marrow studies were not performed. Nevertheless, in our scRNA-seq data we could not identify differences between the analysed pcFCL samples and therefore believe that all our patients had pcFCL. More data is needed to assess whether cutaneous lesions arising from sFCL show different characteristics.

With respect to the staging of primary cutaneous B-cell neoplasms, our data highlights that scRNA-seq can clearly distinguish the investigated subtypes. This is in contrast to sFCL and sDLBCL where previous scRNA-seq studies showed overlapping phenotypes and high inter-patient variability^40,41^. Current recommendations for the diagnostic workup of suspected pcMZL and pcFCL patients requires at least imaging to rule out secondary lesions of systemic lymphoma^11^. While bone marrow studies are no longer recommended for pcMZL, there is currently no consensus on its role in the workup of pcFCL patients^11^. Our data thus holds the promise that scRNA-seq of cutaneous samples may be sufficient to distinguish between primary cutaneous and systemic subtypes of B cell lymphomas arising in the skin.

Finally, our data underlines the recent reclassification of pcMZL as a lymphoproliferative disorder and will need to spark a discussion on the appropriate therapeutic^61,62^ and diagnostic strategies for this indolent disease.

## Acknowledgements

This work was funded by research grants from the Austrian Science Fund to PMB (grant number KLI 849-B) and JG (grant number P35937). SNW was supported by research grants from the Austrian Science Fund (grant number P31127 and IPPTO project number DOC 59-B33). The Vienna Scientific Cluster (Project No. 71839) is gratefully acknowledged for providing computational resources.

## Author Contributions

**Designed research** JG, CJ, SNW, PMB **Sample acquisition** JG, CJ, PMB, SP, WD **Histopathological analysis** MD, ISK **Sample analysis** LS, UM, MF, BA, BML, CW, SNW, WW **Data analysis** JG, SG, IO, VN **Acquisition of funding** JG, PMB **Writing of manuscript** JG, CJ, PMB, SNW, BML

