## Supplementary Materials for "Indolent primary cutaneous B-cell lymphomas resemble persistent antigen reactions without signs of dedifferentiation"

### Indolent primary cutaneous B cell lymphoma represent persistent antigen reactions without signs of dedifferentiation - Supplementary Materials

Johannes Griss MD, PhD[1*], Sabina Gansberger MSc[1], Inigo Oyarzun[1], Mathias C. Drach MD[1], Vy Nguyen MSc[1], Lisa E. Shaw MSc[1], Ulrike Mann[1], Stefanie Porkert MD[1], Matthias Farlik PhD[1], Wolfgang Weninger MD[1], Werner Dolak, MD[2], Bertram Aschenbrenner, PhD[1], Beate M. Lichtenberger, PhD[1], Christine Wagner[1], Ingrid Simonitsch-Klupp MD[3], Stephan N. Wagner MD[1], Constanze Jonak MD[1], Patrick M. Brunner MD, MSc[4*]

1 Department of Dermatology, Medical University of Vienna, Vienna, Austria

2 Division of Gastroenterology and Hepatology, Department of Internal Medicine 3, Medical University of Vienna, Austria

3 Department of Pathology, Medical University of Vienna, Austria

4 Department of Dermatology, Icahn School of Medicine at Mount Sinai, New York, USA

##### Table of Contents

Supplementary Figure 1 2
Supplementary Figure 2 3
Supplementary Figure 3 3
Supplementary Figure 4 5
Supplementary Figure 5 6
Supplementary Figure 6 7
Supplementary Figure 7 8
Supplementary Figure 8 9
Supplementary Figure 9 10
Supplementary Tables 11

### Supplementary Figure 1

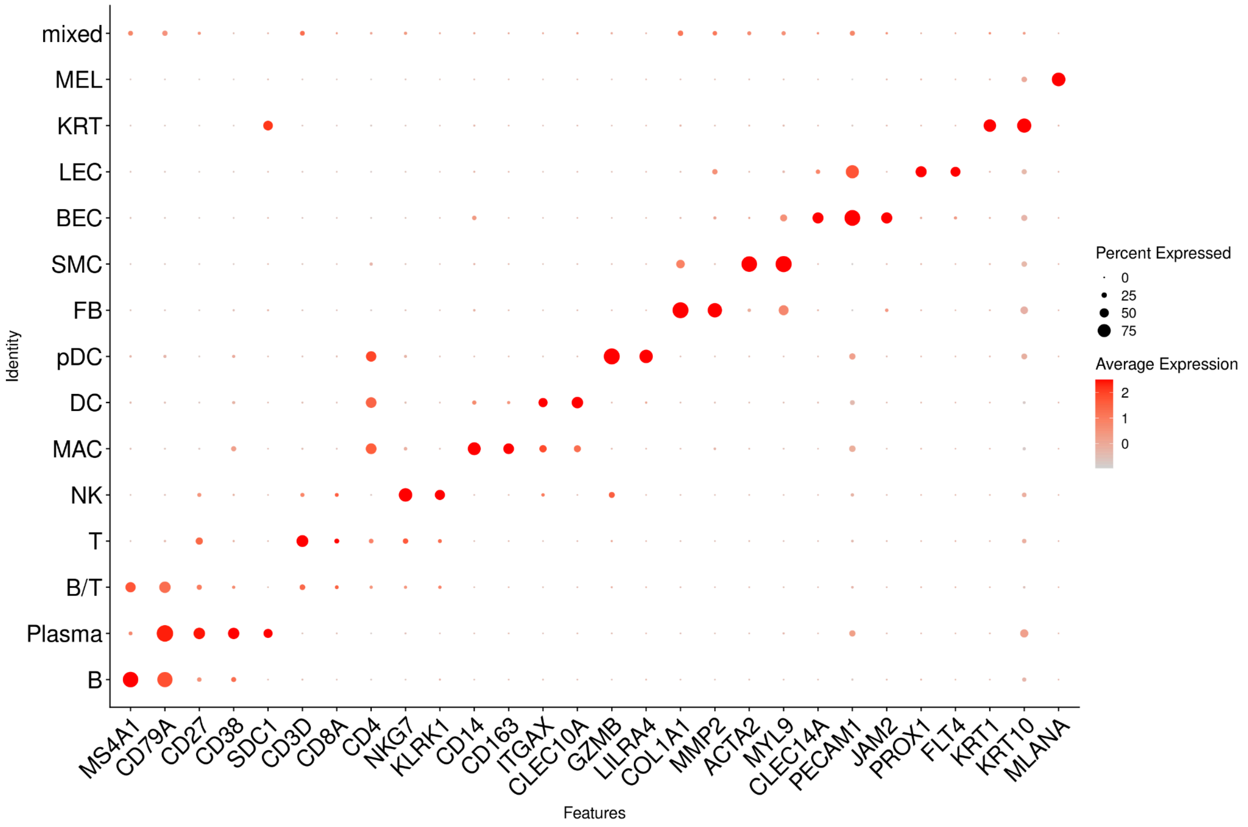


Dot plots of cell clusters from the integration of all cutaneous samples displaying average gene expression (red color) and frequency (circle size) of selected canonical cell type markers.

### Supplementary Figure 2

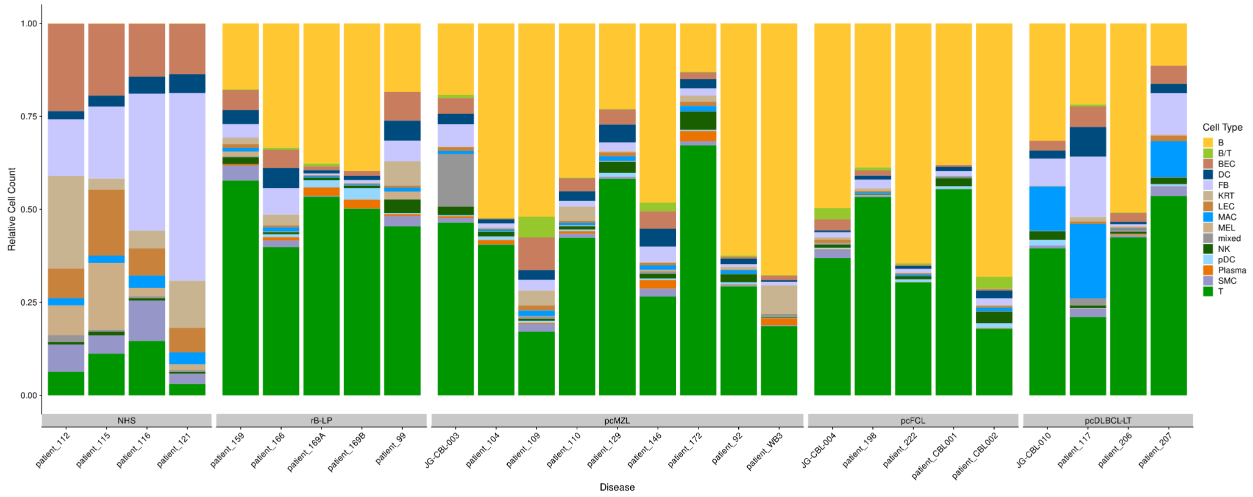


Relative proportion of cell types in each individual cutaneous sample.

### Supplementary Figure 3

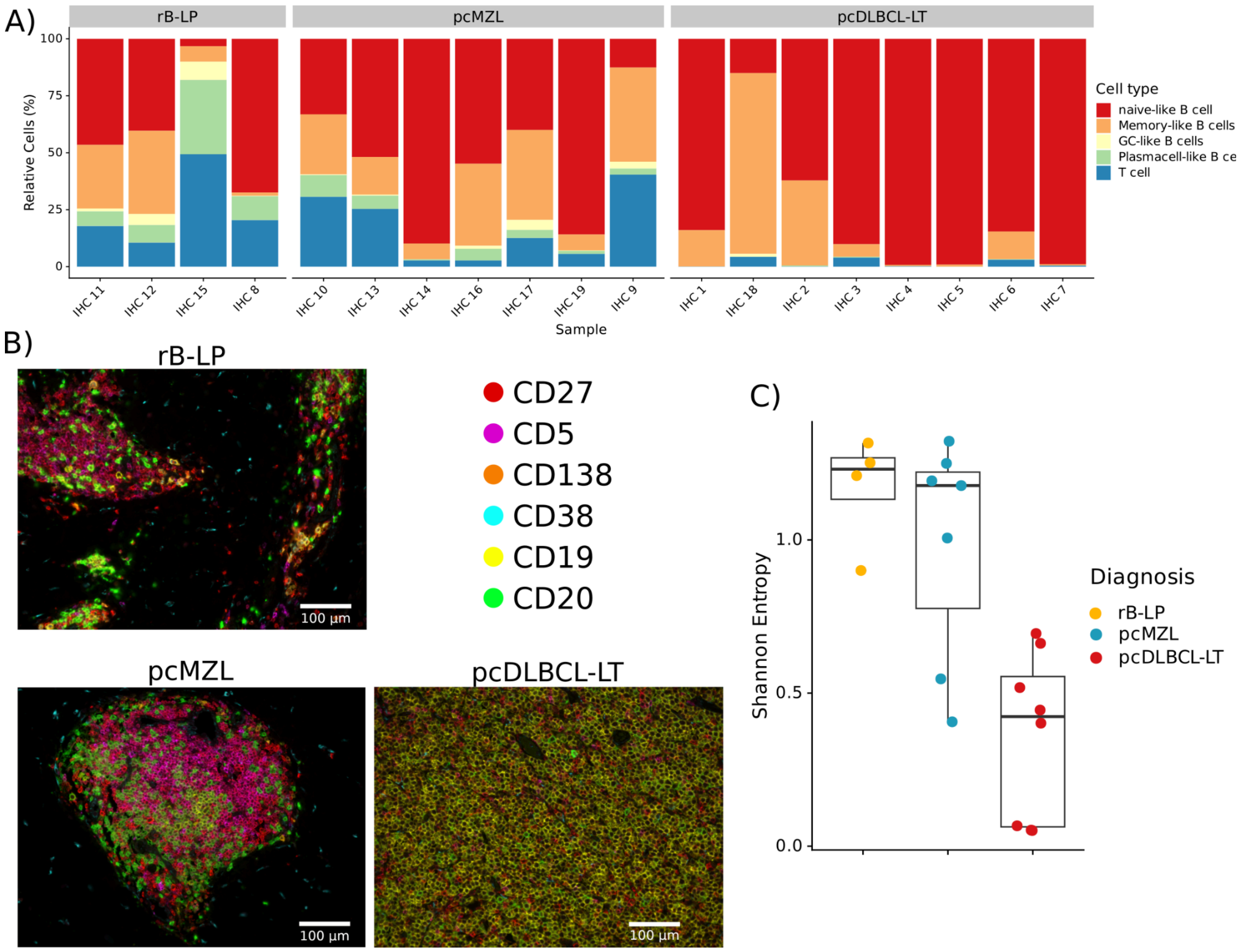


We used a multiplex immunohistochemistry (IHC) approach to simultaneously characterize naive-like B cells (CD19+/CD20+, CD27-, CD38-, CD138-), GC-like B cells (CD19+/CD20+, CD27-, CD38+, CD138-), memory-like B cells (CD19+/CD20+, CD27+, CD38-, CD138-), plasma cells (CD19+, C20-, CD138+), and T cells (CD19-/CD20-, CD5+). Overall, pcMZL and rB-LP samples harbored a diverse lymphocytic infiltrate, as opposed to pcDLBCL-LT matching our scRNA-seq data (A-B). While we observed all B-cell subtypes in rB-LP and pcMZL samples, several pcDLBCL-LT samples only consisted of naive-like CD19+ and CD20+ B cells. A Shannon diversity index based analysis confirmed a significantly larger heterogeneity of lymphocyte composition in rB-LP and pcMZL samples compared to pcDLBCL-LT (Wilcoxon rank sum test, $p < 0.01$,C).

### Supplementary Figure 4

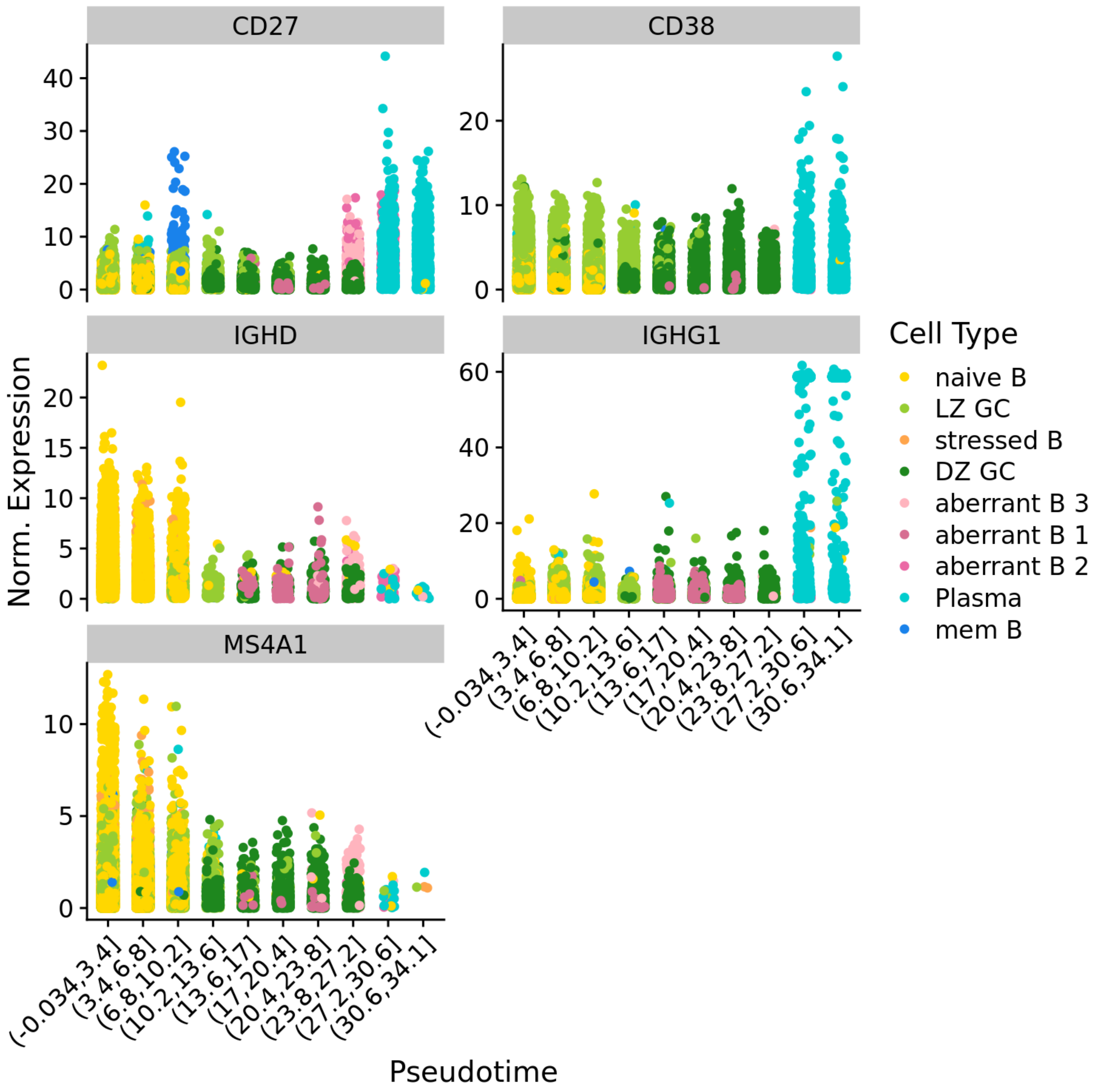


Expression values of key B cell development markers *vs.* the derived pseudotime. Individual points represent individual cells. Colors represent the respective cell phenotypes.

### Supplementary Figure 5

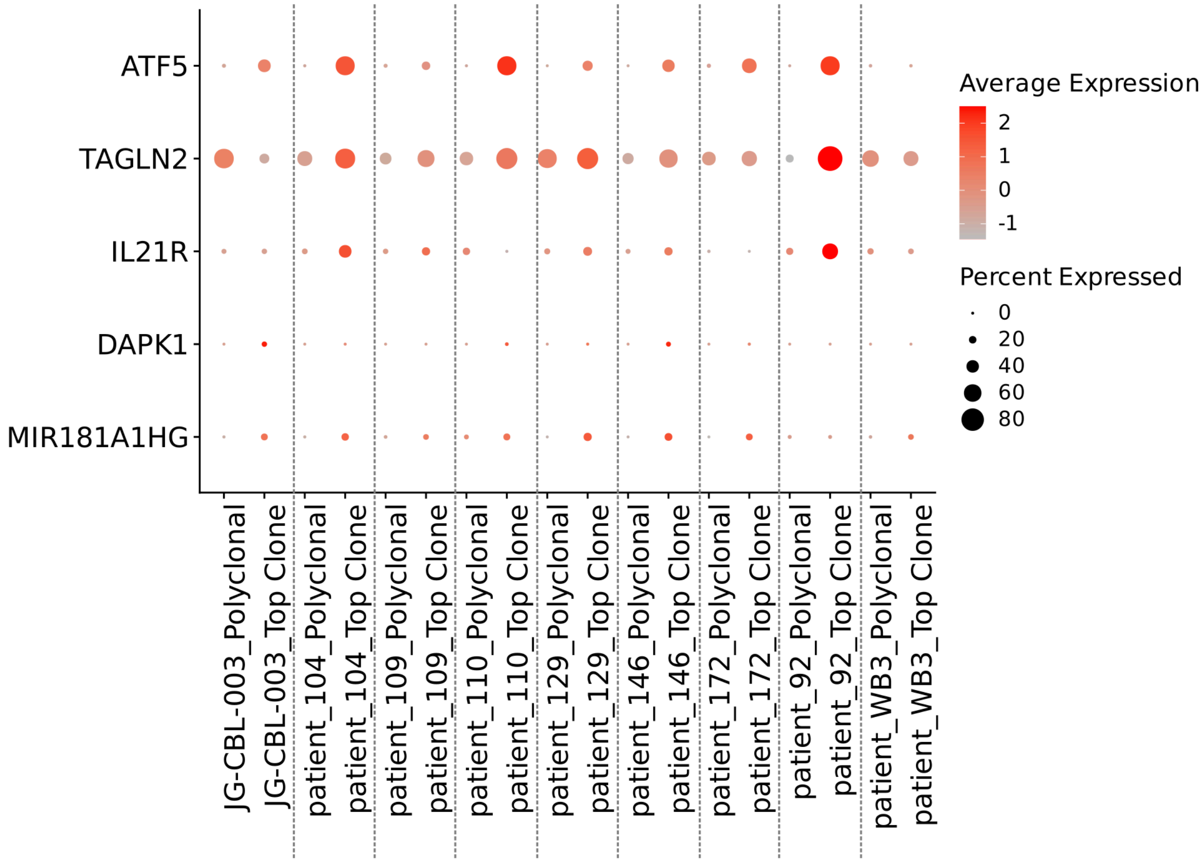


Expression levels of key differentially expressed genes between clonally expanded B cells and polyclonal B cells in pcMZL lesions. Circle sizes represent the fraction of cells expressing the respective genes, color intensity correlates to the expression level.

### Supplementary Figure 6

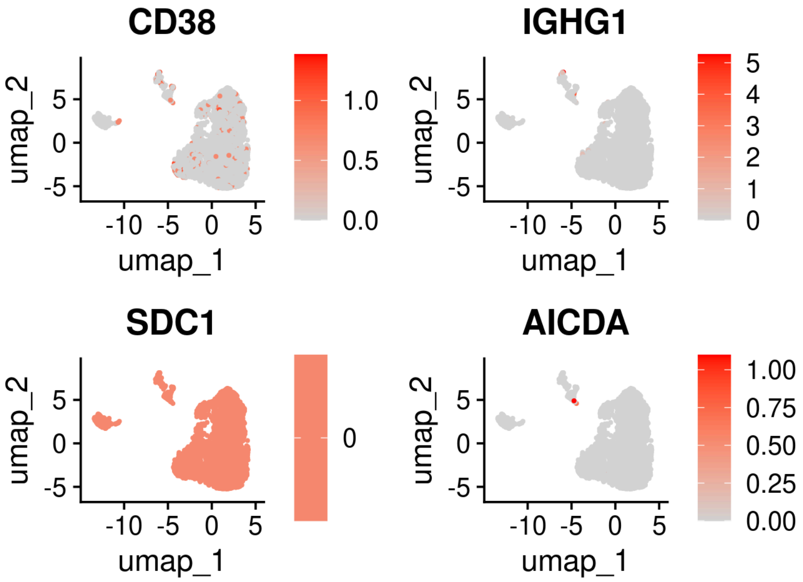


UMAP embedding of a subclustering of B cells from sample 92. Colors represent the expression values of the respective marker genes.

### Supplementary Figure 7

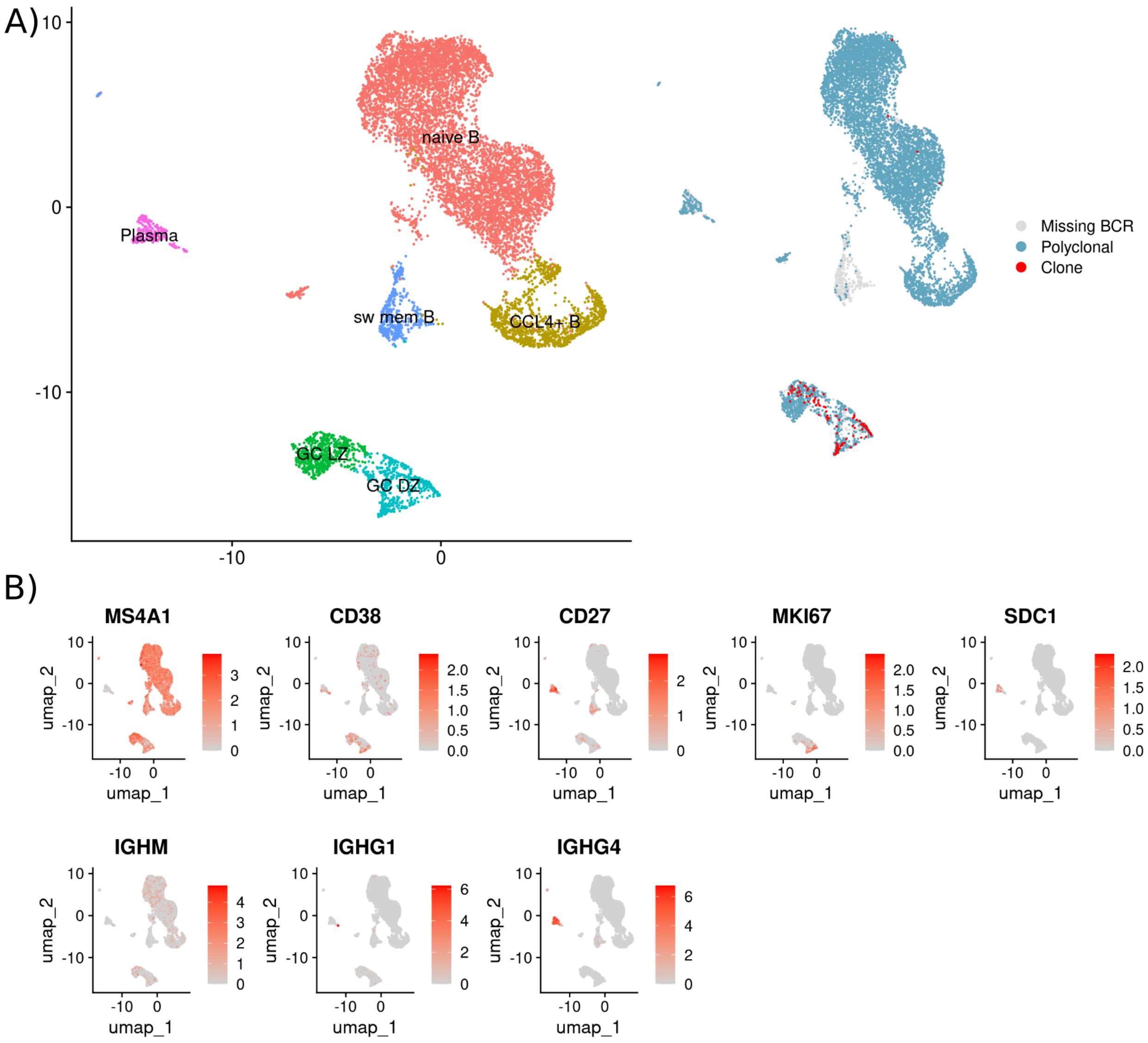


UMAP embedding of a subclustering of B cells from samples WB3. **A)** Colors represent the cell annotation in the left panel, and the BCR information in the right one. **B)** Colors represent the normalized expression values of the respective genes.

### Supplementary Figure 8

A)

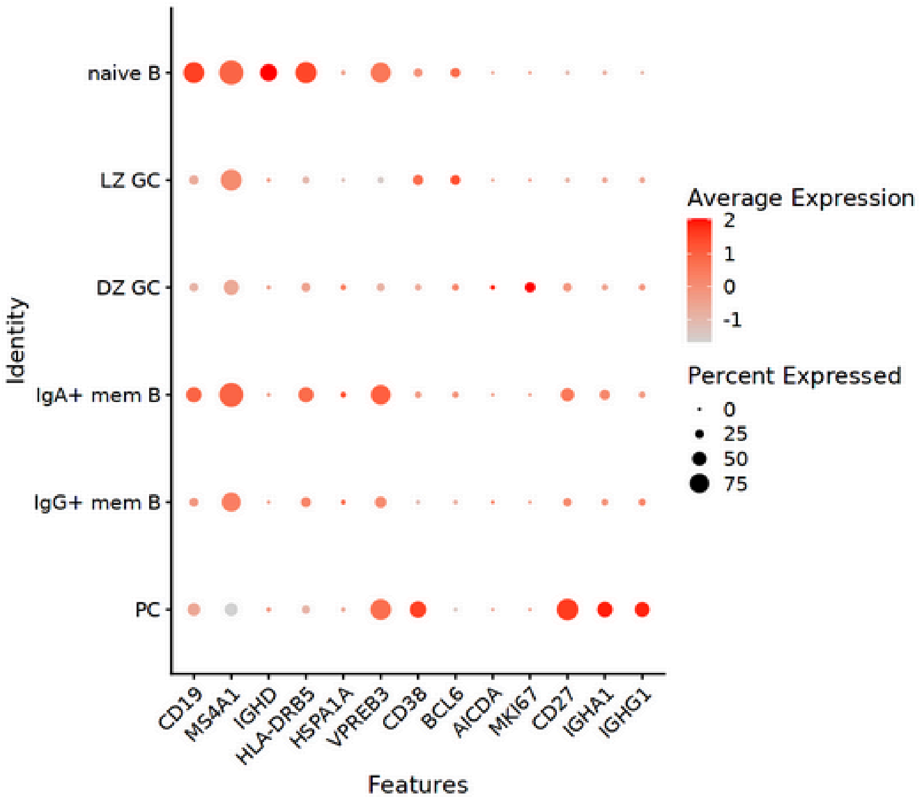


B)

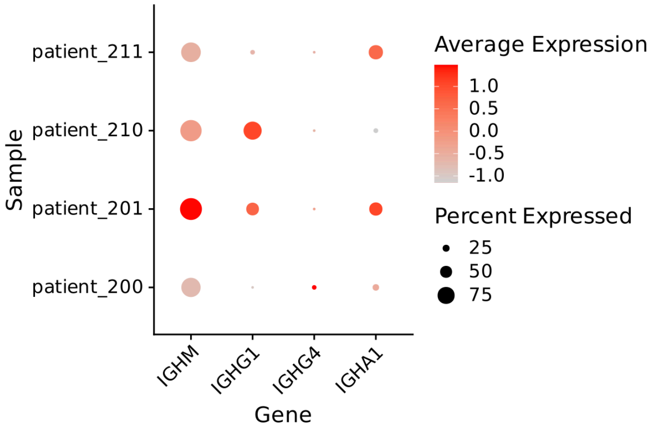


**A)** Dot plots of cell clusters from the subclustering of B cells from the gastric MALT lymphoma samples displaying average gene expression (red color) and frequency (circle size) of selected canonical cell type markers. **B)** Dot plot showing the expression of IG classes for B cells from the top expanded clone per sample.

### Supplementary Figure 9

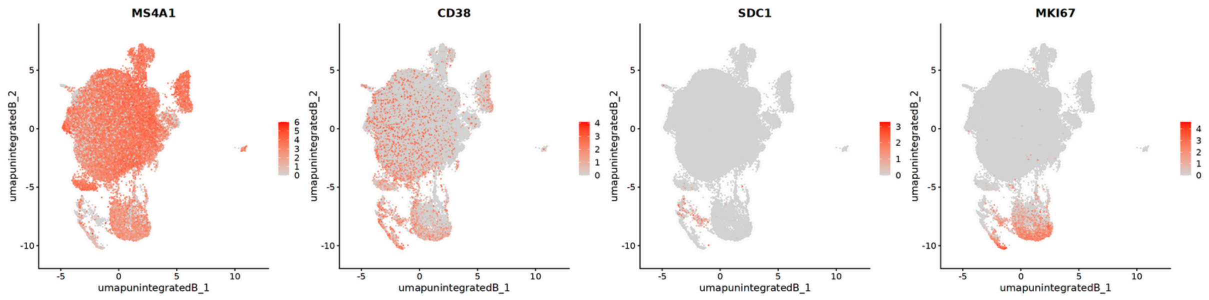


UMAP embedding of the sFCL samples showing the expression values of key B cell markers as color intensities.

### Supplementary Figure 10

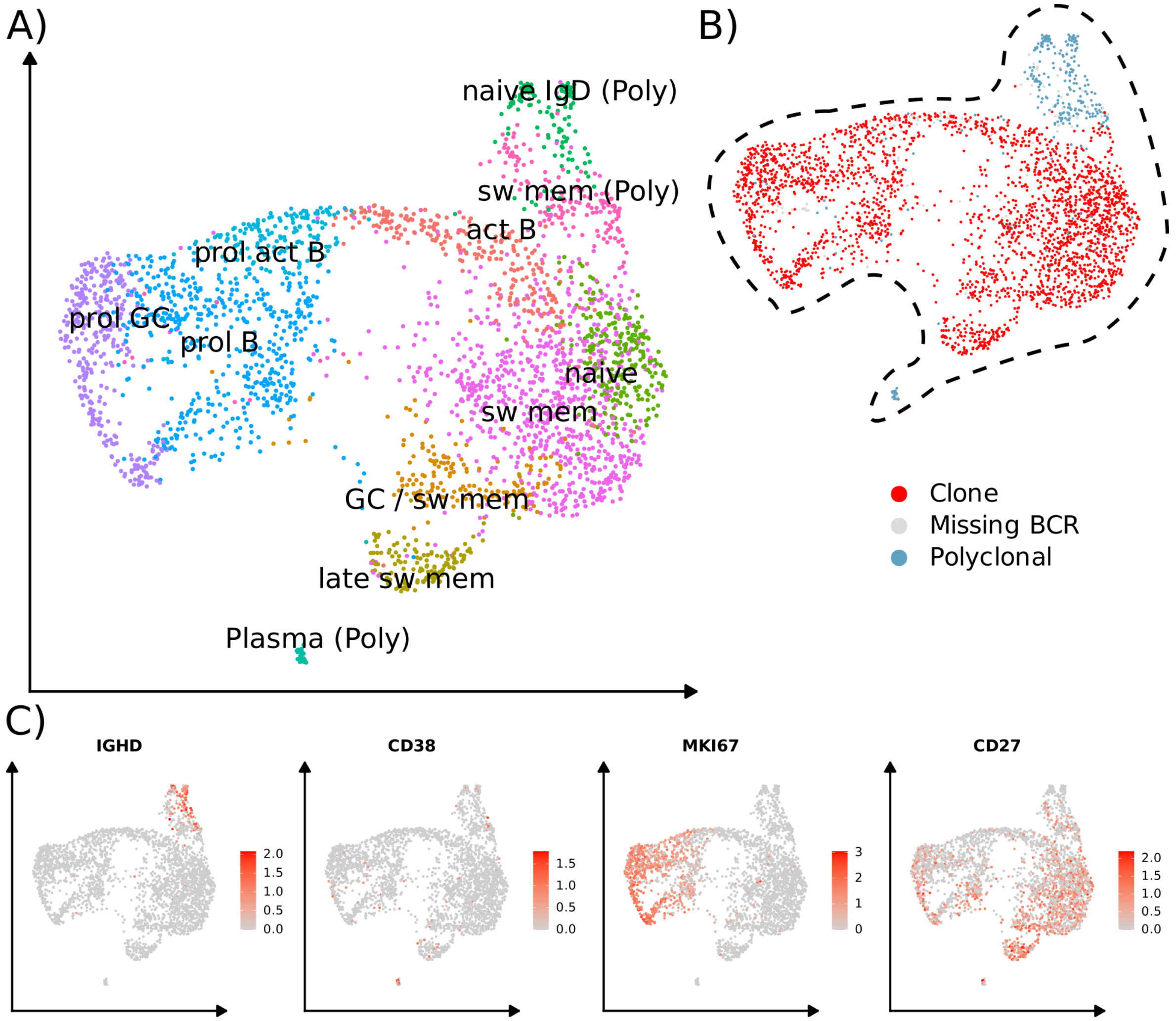


UMAP embedding of the clustering of all B cells from a patient with pcDLBCL-LT from a study on the effect of an oncolytic viral therapy for primary cutaneous B cell lymphoma by Ramelyte et al. A) Colors represent the identified B cell phenotypes. B) Results of the BCR sequencing indicating the top expanded clone and polyclonal B cells. C) Colors represent the expression levels of key B cell markers.

### Supplementary Figure 11

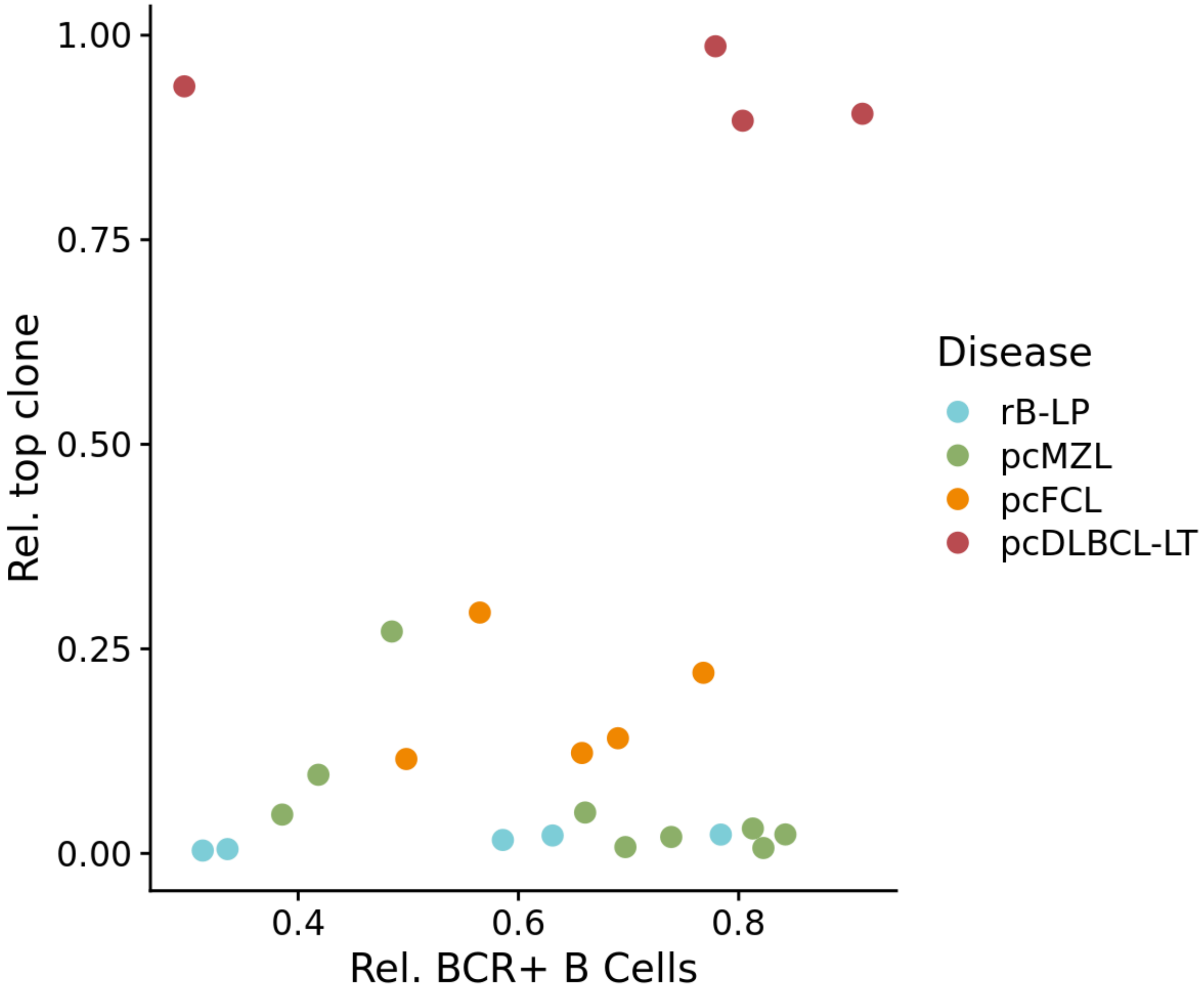


Proportion of B cells with BCR information *vs.* the relative proportion of the top clone of the B cell infiltrates with BCR information for each sample. Color represents the respective diseases. There were three pcMZL samples with a clonal expansion of 9%, 17% and 30%. Yet, these higher rates of clonally expanded B cells correlated with the lowest BCR capture of only 50% of B cells in these samples.

### Supplementary Figure 12

A)

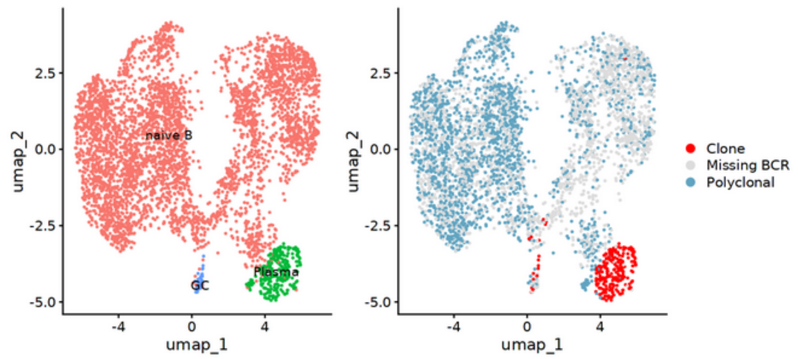


B)

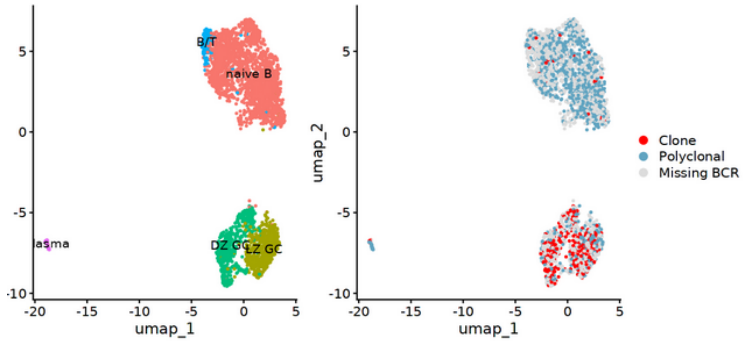


C)

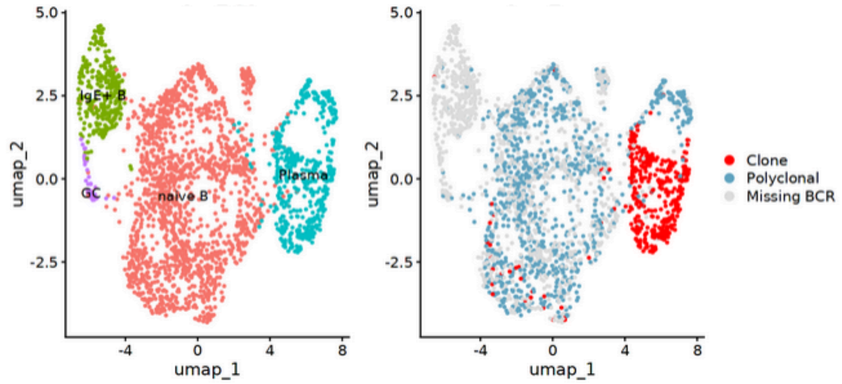


UMAP embedding of a subclustering of the B cells from samples **A)** 146, **B)** 109, **C)** 172. In all samples, B cells without BCR information were found in the naive B cell clusters, indicating that these mainly contained polyclonal B cells. Colors represent the cell annotation in the left panel, and the BCR information in the right one.

### Supplementary Tables

**Supplementary Table 1** Patient baseline characteristics at time of sampling.

| Subject ID | Age | Sex | Race | Histopathological diagnosis | Location | Disease duration (years) | Ongoing treatment | Previous treatments | Disease stage |
|---|---|---|---|---|---|---|---|---|---|
| 112 | 51 | F | White | Healthy control | Trunk | n.a. | n.a. | n.a. | n.a. |
| 115 | 48 | M | White | Healthy control | Trunk | n.a. | n.a. | n.a. | n.a. |
| 116 | 56 | F | White | Healthy control | Trunk | n.a. | n.a. | n.a. | n.a. |
| 121 | 44 | F | White | Healthy control | Trunk | n.a. | n.a. | n.a. | n.a. |
| 92 | 72 | M | White | pcMZL | Upper arm | 8 years | None | Rituximab | T1bN0M0 |
| 104 | 33 | M | White | pcMZL | Trunk | 3 years | None | Clarithromycin | T1bN0M0 |
| 110 | 47 | M | White | pcMZL | Trunk | 13 years | None | Rituximab | T3aN0M0 |
| 129 | 38 | M | White | pcMZL | Shoulder | 14 years | None | Clarithromycin, surgery | T3aN0M0 |
| 146 | 47 | M | White | pcMZL | Upper arm | 1 year | None | Surgery | T1aN0M0 |
| 172 | 72 | W | White | pcMZL | Lower thigh | 2 years | None | Surgery, radiotherapy | T1bN0M0 |
| WB3 | 60 | M | White | pcMZL | Neck | 1 year | None | None | T1bN0M0 |
| 109 | 79 | F | White | pcMZL | Back / Shoulder | 1 year | None | Radiotherapy | T2aN0M0 |
| JG-CBL-003 | 79 | M | White | pcMZL | Back | 3 years | None | Surgery | T1bN0M0 |
| 99 + 159 | 30 | M | White | rB-LP | Upper arms | 1 year | None | None | n.a. |
| 166 | 47 | F | White | rB-LP | Face | 16 years | None | Glucocorticosteroids, hydroxychloroquine, apremilast | n.a. |
| 169A + 169B | 56 | M | White | rB-LP | Upper leg | 1 year | None | Surgery | n.a. |
| 198 | 77 | M | White | pcFCL | Scalp | 1 year | None | None | T2aN0M0 |
| 222 | 32 | M | White | pcFCL | Face | < 1 year | None | Surgery | T2aN0M0 |
| JG_CBL_001 | 51 | M | White | pcFCL | Scalp | 3 years | None | None | T2cN0M0 |
| JG_CBL_002 | 72 | W | White | pcFCL | Face | < 1 year | None | None | T2cN0M0 |
| JG-CBL-004 | 43 | M | White | pcFCL | Scalp | 10 years | None | Rituximab | T2cN0M0 |
| 206 | 74 | M | White | pcDLBCL-LT | Lower leg | <1 year | None | None | T2bN0M0 |
| 117 + 207 | 78 | F | White | pcDLBCL-LT | Forearm and lower leg | <1 year | None | Radiotherapy, R-CHOP | T3aN0M0 |
| JG-CBL-010 | 79 | F | White | pcDLBCL-LT | Lower leg | <1 year | None | None | T2bN0M0 |
| 200 | 62 | M | White | Gastric MALT | -- | < 1 year | None | HP-Eradication | n.a. |
| 201 | 31 | M | White | Gastric MALT | -- | 1 year | None | HP-Eradication | n.a. |
| 210 | 59 | F | White | Gastric MALT | -- | 2 years | None | HP-Eradication | n.a. |
| 211 | 63 | F | White | Gastric MALT | -- | 1 year | None | HP-Eradication | n.a. |
